# Assessment of *in vivo* epithelial disruption to model anorectal *Chlamydia* infection in mice

**DOI:** 10.64898/2026.10.24.761026

**Authors:** Zoe E.R. Wilton, Susan B. Core, Leslie M. Huggins, Andzoa N Jamus, Isabella G. Romano, James P. Davis, Anthony D. Cretara, Kathryn M. Frietze

## Abstract

*Chlamydia trachomatis* (Ct) causes the most commonly diagnosed bacterial sexually transmitted infection. Ct can infect mucosal epithelial cells throughout the human body and is frequently found at the anorectal tract. However, there are currently few models of anorectal *Chlamydia* infection, limiting our ability to investigate vaccines, identify therapeutics, and understand infection dynamics. Since epithelial disruption prior to infection has been used in other STI animal models to increase permissibility and improve reproducibility of infection, we evaluated the impact of different methods of epithelial disruption prior to inoculation on the bacterial burden of anorectal *Chlamydia muridarum* (Cm). Mice were treated anorectally with physical (calcium alginate swab) and/or chemical (nonoxynal-9) disruption 24 hours prior to anorectal inoculation with 1 x 10^4^ IFUs of a luciferase-expressing Cm (Luc-Cm). Infection progression was monitored in real time using an *in vivo* imaging system (IVIS) and the shedding of Luc-Cm DNA in the stool was quantified by qPCR. Cm was detected in all infected treatment groups by both IVIS and qPCR, irrespective of treatment. However, N-9 treatment prior to inoculation with Luc-Cm increased infection reproducibility and persistence when monitored using IVIS imaging, while having minimal adverse reactions. Therefore, chemical disruption prior to Luc-Cm infection was utilized to examine the efficacy of a bacteriophage virus-like particle (VLP) vaccine against Luc-Cm (Qβ-VD4-Cm) in the anorectal tract. We found that immunization with Qβ-VD4-Cm did not reduce Luc-Cm infection in the anorectal tract, despite this vaccine previously being shown to reduce Luc-Cm infection in the female reproductive tract. This highlights the unique nature of vaccine-mediated protection that can occur at distinct anatomical sites of infection with the same pathogen.

## 1. INTRODUCTION

*Chlamydia trachomatis* (Ct) is the most commonly diagnosed cause of bacterial sexually transmitted infections (STIs) globally, with 129 million newly diagnosed cases reported in 2020 [1,2]. Ct infects mucosal epithelial cells, with sexually transmitted Ct often detected at urogenital, anorectal, and oropharyngeal sites [3,4]. Most research on Ct has focused on female reproductive tract (FRT) infection due to the associated significant pathological sequelae and increased risk of infertility. However, anorectal infections are common in males and females [4]. Anorectal Ct, although frequently asymptomatic, can cause a myriad of sequelae including fever, non-specific inflammation, rectal pain, proctitis, colitis, anal ulcers, fistulas, rectal abscesses, and granulomas [4,5]. Anorectal Ct infections have also been associated with an increased risk of HIV acquisition in men who have sex with men (MSM) [6,7]. In addition, anorectal Ct infections have been hypothesized to be a reservoir for autoinoculation and reinfection of the urogenital tract, particularly in women [4,8,9]. Conversely, anorectal Ct infections are hypothesized to promote the development of protective immune responses against urogenital Ct infections [8,10,11]. This is supported by experiments showing that rectal administration of a live-attenuated *Chlamydia* vaccine resulted in reduced pathology in the FRT of mice, increased serum IgG, fecal IgA, and T cell responses in infected mice [10]. This has important implications for vaccine development and antibiotic treatment efficacy.

*Chlamydia muridarum* (Cm) is a well-studied mouse pathogen that is used to model key features of urogenital *Chlamydia* infections [12,13]. Intravaginal inoculation with Cm recapitulates key features of human Ct FRT infections, resulting in ascension of the pathogen into the upper reproductive tract and hydrosalpinx in mice [14,15]. However, while intravaginal Cm infection is a well-established and characterized model, anorectal Cm infection models are less well-established. Anorectal inoculation of mice with Cm has been previously reported [8,16]. These studies show that persistent infection can be established via direct inoculation of Cm into the anorectal tract, resulting in high bacterial burden early during infection that tapers to low-level, persistent infections [8,16]. One model, established in Slade et al., rectally infected 8–10-week-old nectin-1 knockout male mice (B6.129X1(Cg)-Pvrl1tm1Ytk/J) with 1 x 10^6^ IFU of Cm using a catheter that was placed 3 cm into the rectal tract before inoculation [16]. Another model, established in Wang et al., used a pipette tip to directly instill 5 x 10^4^ or 1 x 10^7^ IFUs of Cm into the rectum of 6–7-week-old female mice [8]. This experiment utilized several strains of female mice for rectal infection: CBA/J, C57BL/6J, and C3H/HeJ. Both models were successfully able to establish Cm infection in the rectal tract of mice. Recognizing that the amount of infectious inoculum used in these models is orders of magnitude more than would be expected in natural infection, we sought to develop an anorectal mouse model of *Chlamydia* infection that would be relevant for vaccine testing.

Chemical or physical epithelial disruption prior to inoculation is a common technique to increase reproducibility and permissibility of mice for sexually transmitted pathogens. For instance, mouse models of Human Papillomavirus (HPV) have shown that epithelial cell disruption prior to inoculation of either HPV16 pseudovirus or mouse papillomavirus MmuPV1 increases early viral load in both the urogenital and anorectal tracts of BALB/cAnNCr and *Foxn1^nu^*/*Foxn1^nu^* mice, respectively [17,18]. Here, we investigated the use of chemical and/or physical disruption of the anorectal epithelium prior to Cm infection as a method for increasing the permissibility and reproducibility of this model. We also used this model to test the efficacy of a candidate vaccine to protect against Cm anorectal infection.

## 2. MATERIALS AND METHODS

### 2.1. *Chlamydia muridarum* propagation and isolation

Luciferase expressing *Chlamydia muridarum* (Luc-Cm, which was a generous gift from Dr. Guangming Zhong at the University of Texas Health Science Center in San Antonio) was propagated in McCoy cells (CRL-1696, ATCC, Manassas, VA, USA). Briefly, Luc-Cm was released from the McCoy cells via physical disruption using sterile glass beads and sonication. Elementary bodies (EBs) were then isolated using a 30% Gastrografin (Bracco Diagnostics, Monroe Twp., NJ, USA) density gradient solution and resuspended in medium containing 250 mM sucrose, 10 mM sodium phosphate, and 5 mM L-glutamic acid (SPG buffer). The isolated EBs were stored at −80°C until use. To quantify the inclusion forming units (IFU) present, the EBs were titered on McCoy cell monolayers and counted using a fluorescence microscope[19].

### 2.2 Ethics Statement for Animal Studies

All animal studies were reviewed and approved by the Institutional Animal Care and Use Committee (IACUC) of the University of New Mexico (UNM) School of Medicine (Protocol number 25-201630). Animals were housed in the Animal Research Facility at UNM and maintained following IACUC guidelines. Weight loss (reported as precent weight change) and diarrhea were monitored throughout the study. Mice that experienced excessive weight loss (greater than 20% of the original body weight) were humanely euthanized.

### 2.3. Mouse Models of Anorectal Infection

6- to 8-week-old BALB/c mice (n= 20: male and females) were purchased from Jackson Laboratories (Bar Harbor, ME, USA). Physical disruption, chemical disruption, both, or no treatment were performed one day prior to anorectal infection with Luc-Cm (Day −1)[14]. For the physical disruption group, Puritan Calgiswab Sterile Nasopharyngeal Calcium Alginate tipped applicators (25-800 A 50) were gently inserted into the anorectal tract of the mice and slowly rotated back and forth 20 times [17]. For the chemical disruption group, a positive displacement pipette was used to instill 20 μL of 4% nonoxynol-9 (N-9) in 3% Carboxymethylcellulose (CMC) into the anorectal tract of the mice [17]. For the physical/chemical disruption group, Puritan Calgiswab Sterile Nasopharyngeal Calcium Alginate tipped applicators were gently inserted into the anorectal tract of the mice and slowly rotated back and forth 20 times. After the applicator was removed, a positive displacement pipette was used to insert 20 μL of 4% N-9 in 3% CMC into the anorectal tract of the mice. On day 0, the no disruption, physical disruption, chemical disruption, and physical/chemical disruption groups were infected with 1 X 10^4^ IFUs of Luc-Cm in 20 μL total volume inoculum delivered with a 20 μL pipette. The negative control group was inoculated with 20 μL of sucrose phosphate glutamate buffer.

### 2.4. Fecal DNA extraction

Fecal samples (<3 pellets) were collected from each mouse on days 1-7, 10, 15, and 20 post-infection. Mice were placed on a sterile surface, and fecal samples were collected with tweezers immediately after defecation. Collected fecal samples were stored at −20°C prior to DNA extraction. DNA was extracted from mouse fecal pellets using the PureLink™ Microbiome DNA Purification Kit (Invitrogen, Carlsbad, CA, USA), following the manufacturer’s protocol. Eluted DNA was stored at −20°C until use.

### 2.5. Real-Time qPCR

The amount of Luc-Cm DNA extracted from fecal samples was determined using Real-Time qPCR [20]. Real-Time qPCR reactions to detect *Chlamydia* 16S included 6.25 mL of TaqMan Fast (Thermo Fisher Scientific, Waltham, MA, USA), 0.35 mL of 10μM Forward 16S Primer (5’-GGA GGC TGC AGT CGA GAA TCT −3’: Integrated DNA Technologies (IDT), Coralville, IA, USA), 0.35 mL of 10μM Reverse 16S Primer (5’-TTA CAA CCC TAG AGC CTT CAT CAC A −3’: IDT), 0.2 mL of 25μM 16S Probe (5’-/56-FAM/TCG TCA GAC TTC CGT CCA TTG CGA /36-TAMSp/ −3’: IDT), 2.85 mL of RNase free water and 10 μL of extracted fecal DNA per reaction. Each sample was run in duplicate. Reactions carried out using the Applied Biosystems QuantStudio 7 Flex Real-Time PCR System (Thermo Fisher Scientific) using the following cycling conditions: hold at 50°C for 120 seconds, hold at 95°C for 20 seconds, and then a 2-step amplification was performed for 40 cycles that consisted of running at 95°C for 1 second and then 60°C for 20 seconds. The picograms of Luc-Cm DNA was then determined using the mean Ct value of each sample and a standard curve generated from Luc-Cm DNA.

### 2.6. Cm EB ELISA

EB ELISA was performed as previously described [1]. Briefly, Immulon 2 HB 96-well flat-bottom microtiter plates (Thermo Scientific, Waltham, MA, USA) were coated with 75 μL/well of 1 μg/mL poly-L-lysine (diluted in 0.05 M bicarbonate buffer: pH 9.6) and incubated overnight at 4°C. The plates were then washed three times with 0.05% PBS–Tween-20 (PBST). After washing, 50 μL of Cm EBs (4.5 μg total protein/mL, in PBS) was added to each of the wells. After Cm EBs were added, the plates were centrifuged at 900 x g for 5 mins at room temperature and then fixed for 5 mins at room temperature using 50 μL/well of 0.1% glutaraldehyde in PBS. Plates were then washed 3 times with PBST followed by blocking with 300 μL/well of 2% goat serum-PBST and overnight incubation at 4°C. Plates were then decanted and 100 μL/ well of serially diluted immune sera (in 2% goat serum-PBST) was added. Plates were incubated at room temperature for 2 hours with rocking followed by washing four times with PBST. After washing, 100 μL/well of either peroxidase-conjugated anti-mouse IgG (Jackson ImmunoResearch, West Grove, PA, USA) or IgA (SouthernBiotech, Birmingham, AL, USA) secondary antibody (diluted 1:5000 in 2% goat serum-PBST) was added and incubated at room temperature for 30 mins with rocking. Plates were then washed four times with PBST. After washing, 100 μL/well of 3,3′,5,5′-tetramethylbenzidine (TMB: EMD Millipore, Burlington, MA, USA) was added to the plates and incubated at room temperature for 20 mins with rocking. After incubation, the plates were immediately quenched with 1% hydrochloric acid and the absorbance was read at 450 nm using accuSkan FC plate reader (Fisher Scientific, Waltham, MA, USA). The plate background reading (empty wells) was subtracted from the well values.

### 2.7. VD4 Peptide ELISA

VD4 peptide ELISA was performed as previously described [1]. Briefly, Immulon 2 HB 96-well flat-bottom microtiter plates were coated with 50 μL/well of 10 μg/mL streptavidin (diluted in PBS: Invitrogen, Carlsbad, CA, USA) and incubated overnight at 4°C. The plates were then washed three times with PBS. After washing, 50 μL of 10 μg/mL Succinimidyl 6-(beta- maleimidopro pionamido)-hexanoate (SMPH: Thermo Fisher Scientific, Waltham, MA, USA), diluted in PBS was added to each of the wells and the plates were incubated for 1 hour at room temperature with rocking. Plates were then washed 3 times with PBS and 1 μg/50 μL of the VD4 peptide was added to each well. After incubating at room temperature for 2 hours with rocking, the plates were washed with PBS three times and blocked with 150 μL/well of 0.5% milk in PBS. After blocking overnight at 4°C, the plates were washed three times with PBS and 50 μL/well of serially diluted immune sera (in 0.5% milk in PBS) was added. Plates were incubated at room temperature for 2 hours with rocking followed by washing five times with PBS. After washing, 50 μL/well of either peroxidase-conjugated anti-mouse IgG (Jackson ImmunoResearch, West Grove, PA, USA) or IgA (SouthernBiotech, Birmingham, AL, USA) secondary antibody (diluted 1:5000 in 0.5% milk in PBS) was added and the plates were incubated at room temperature for 45 minutes with rocking. Plates were then washed three times with PBS. After washing, 50 μL/well of TMB was added to the plates and incubated at room temperature for 15 (IgG secondary antibody) or 25 mins (IgA secondary antibody) with rocking. After incubation, the plates were immediately quenched with 50 μL/well of 1% hydrochloric acid and the absorbance was read at 450 nm using accuSkan FC plate reader.

### 2.8. IVIS imaging

As previously described, IVIS (PerkinElmer, Waltham, MA, USA) was used post infection (days 1-7, 10, 15, and 20) to examine Luc-Cm infection [1]. D-Luciferin (20 μL of 20 mg/mL, diluted in PBS: Perkin Elmer, Shelton, CT, USA) was instilled into the anorectum of the mice using a 20 μL pipette. The mice were then anesthetized using inhaled isoflurane and three minutes post D-luciferin instillation, images were acquired with a 5 min exposure. The average radiance (p/s/cm2/sr) was determined by selecting uniform regions of interest (ROIs) on the abdomen of each mouse. Background average radiance was determined by subtracting the average radiance of the ROIs from the uninfected mice from experimental subjects. Data were collected using Live Image 4.3.1 software (Revvity, Waltham, MA, USA).

### 2.9. Production of Bacteriophage Qβ VLPs

Bacteriophage Qβ VLPs were produced in *Escherichia coli* (E. coli) as previously described. Briefly, E. coli C41 electrocompetent cells (Sigma Aldrich, Burlington, MA, USA) were transformed with a plasmid that encodes for the Qβ coat protein and a Kanamycin resistance gene, which allows for antibiotic-mediated selection. Qβ-expressing E. coli were grown in 500 mL of LB broth until the culture reached an optical density (O.D.) of 0.6-0.8, at which time 0.5 mM isopropyl β-D-1-thiogalactopyranoside (IPTG: Sigma Aldrich, Burlington, MA, USA) was added to induce Qβ coat protein production. After incubating overnight at room temperature, the E. coli cultures were centrifugated to remove the supernatant, and the pellets were stored at −80°C. To lyse the E. coli cells and isolate the Qβ coat protein, the pellets were thawed and then incubated with 100 mL of lysozyme solution (100 mM NaCl, 10 mM EDTA, and 50 mM Tris-HCl dissolved in diH2O) and 0.5 mL of 10% deoxycholate (DOC) for 1 hour at 4°C. After incubation, the pellets were briefly sonicated (Qsonica, Newtown, CT, USA). Then 100 μL of 10 mg/mL DNase and 2.5 mL of 2 M MgCl2 was added to the pellet solution and incubated for 1 hour at 37°C. After incubation, the pellet solution was centrifuged and the supernatant was collected. Ammonium sulfate was then added at 60% saturation and incubated overnight at 4°C with rocking. The solution was then centrifuged at 15,317 x g for 10 mins and the supernatant was discarded. The pellet was then resuspended in 1x SCB buffer (100 mM NaCl, 10 mM Tris-HCl, and 2.5 mM MgSO_4_ dissolved in diH2O), centrifugated, and the supernatant was collected. This process was repeated one more time and then the combined supernatant was stored at −80°C until size-exclusion chromatography.

For size exclusion chromatography, the supernatant was thawed and then centrifuged. After centrifugation, the supernatant was collected and loaded on a CL-2B Sepharose column and separated by gravity flow with SCB buffer overnight at 4°C with fractionation. Fractions containing the Qβ VLP were then identified using a 1% agarose gel followed by an SDS-Page gel to identify fractions with VLPs. The combined VLP-containing fractions were then precipitated by 70% saturation ammonium sulfate overnight at 4°C with rocking. The solution was then centrifuged for 10 mins at 15,317 x g and the supernatant was discarded. The pellet was then resuspended in 1x PBS and dialyzed against PBS using 10K MWCO Snakeskin Dialysis Tubing (Thermo Fisher Scientific, Waltham, MA, USA). Endotoxin was then depleted from VLPs by 5 sequential phase extractions with Triton-X 114. Once completed, the purified Qβ VLP was quantified by SDS-PAGE with Coomassie staining and stored at −20°C.

### 2.10. Preparation of Qβ-VD4-Cm VLP vaccine

Purified Qβ VLP was thawed and incubated with 10 molar excess SMPH at room temperature for 2 hours with rocking. Unconjugated SMPH was removed using 100,000 NMWL Amicon Ultra Centrifugal Filter tubes (Merck Millipore, Burlington, MA, USA). Qβ-SMPH was washed with PBS 2 times and then the supernatant containing the Qβ-SMPH was placed in a 1.5 mL tube. 10 molar excess of the Cm VD4 peptide (GenScript, Piscataway, NJ, USA) was then added to the 1.5 mL tube and the solution was incubated overnight at 4°C with rocking. Unconjugated peptide was removed using 100,000 NMWL Amicon Ultra Centrifugal Filter tubes. Two washes with PBS were performed and then the supernatant containing the Qβ-VD4-Cm VLP was collected. Successful conjugation was confirmed using SDS-PAGE and diluted to 5 μg of VLP/ 50 μL with PBS. The VLP vaccines were stored at −20°C until use.

### 2.11. Anorectal Luc-Cm Challenge of VLP Immunized Mice

Male and female BALB/c mice (n = 20/group) were immunized with their respective vaccine (Qβ-VD4-Cm, Qβ, or PBS) 3 times, 3 weeks apart, via intramuscular (I.M.) injection without exogenous adjuvant. Immune sera and fecal pellets were collected every 3 weeks for ELISA to assess antibody titers. Fecal material was prepared for ELISA as previously described [21,22]. Briefly, the fecal pellets were resuspended in 10% Goat Serum in PBST (0.05% Tween 20) at 1:10 *w/v* for an hour with vortexing. Once the pellets were dissolved, the solution was centrifuged and the supernatant was collected for storage at −20°C until use.

On day 68 after first immunization, the mice received chemical disruption treatment in the anorectal tract. The next day 1 x 10^4^ IFUs of Luc-Cm (experimental groups) or SPG (uninfected control group) was inoculated into the anorectal tract. Early bacterial burden was then monitored using IVIS imaging after direct instillation of D-Luciferin (20 μL of 20 mg/mL) into the anorectal tract until day 20 post-infection as previously described. Mice were weighed on all days of the experiment.

### 2.12 Impact of Chemical Disruption: Histology

Male and female BALB/c mice received mock I.M. immunizations with PBS 3 times, 3 weeks apart. On day 68 post-1^st^ immunization, the mice received either no disruption (n=20) or chemical disruption (n=20) with 4% N-9 treatment in the anorectal tract. After 24 hours, fecal material was collected, the mice were euthanized, and the distal colon was collected for histology. 24 hours after epithelial disruption treatment, the mice were euthanized and a 6 cm tissue section of the distal colon was removed. For Hematoxylin and Eosin (H&E) staining, the tissue samples (n=20) were placed in room temperature formalin (Epredia, Portsmouth, NH, USA) for 48 hours. The formalin was then removed, the tissue samples were washed 3 times with room temperature 70% ethanol, and then tissue samples were stored in a 15 mL tube with room temperature 70% ethanol until the samples could be processed. For immunohistochemistry (IHC), the tissue samples (n=20) were immersed in cold Carnoy’s solution (100 mL of Carnoy’s is made with 60 mL absolute ethanol, 30 mL chloroform, and 10 mL glacial acetic acid) for 2 hours. The Carnoy’s solution was then removed, the tissue samples were washed 3 times with cold 100% ethanol, and then tissue samples were stored at −20°C in a 15 mL tube with 100% ethanol until the samples could be processed. The tissue repository core at UNM then processed the samples by paraffin embedding and sectioning the tissue (4-5 μm). For IHC, the tissue was stained using a rabbit anti-MUC2 monoclonal primary antibody (Abcam, Cambridge, United Kingdom) and an anti-rabbit secondary antibody that was conjugated to DAB. Tissue samples were observed by the Leica Aperio AT2 microscopy (Leica Biosystems, Deer Park, IL, USA) for H&E staining and the Ventana Discovery Ulta platform (Ventana Medical Systems, Tucson, AZ, USA) for IHC staining. The slides were also digitized for analysis. For IHC, image analysis and H-scoring was performed using the HALO image analysis platform from IndicaLab (Albuquerque, NM, USA). For H&E, pathology scores were determined by blinded analysis from an intestinal pathologist using a previously described scoring criteria[23].

### 2.13 Fecal LCN-2 and Total Fecal IgA ELISAs

The fecal samples were collected 24 hours post-epithelial disruption treatment, immediately placed in liquid nitrogen for snap freezing, and then stored at −80°C until use. The fecal pellets were then homogenized at 90 mg/ml in 0.1% PBST using a vortex mixer (Greiner Bio-One, Monroe, NC, USA). Once homogenized, the tubes were centrifuged at 20,000 xg for 10 minutes at 4°C. The supernatant was then removed, placed in a clean 1.5 mL Eppendorf tube, and stored at −80°C until use. The LCN-2 (R&D Systems, Minneapolis, MN, USA) and Total IgA (Invitrogen, Carlsbad, CA, USA) ELISAs were performed following the manufacturer’s kit protocols. Fecal samples were thawed and plated directly on the LCN-2 ELISA plate without diluting or on the IgA ELISA plate at a 2-fold dilution. Standards were provided by both kits to generate a known standard curve. The line of best fit was calculated for each ELISA based on the standard curve and used to determine the amount of LCN-2 and Total IgA present in fecal samples.

### 2.14. Statistical Analysis

All of the results were analyzed in GraphPad Prism Version 10.5.0 (673).

## 3. RESULTS

### 3.1 Epithelial disruption prior to Luc-Cm inoculation of the anorectum alters time-course of infection in mice

To investigate alterations to Luc-Cm infection after chemical or physical disruption to the anorectal epithelium, BALB/c mice were treated with nonoxynol-9 (N-9) for chemical disruption, swab for physical disruption, or both prior to inoculation with Luc-Cm. Infection kinetics were followed longitudinally in each mouse by imaging with IVIS on days 1-7, 10, 15, and 20 post inoculation. Males and females were analyzed separately and together (Fig. 1). Overall, males tended to have lower overall infection over the course of the experiment compared to females (Supplementary Fig. 1), although the overall trend of infection in treatment groups was similar (Fig. 1D-F). No statistical difference in overall infection was observed between infected treatment groups (Fig. 1A-C). However, we did observe a statistically significant increase in the number of days of infection for N-9 and swab/N-9 treatment groups compared to the no treatment group for females (7.7 days and 8.0 days respectively vs 4.6 days) (Table 1). We also observed a smaller range of positive days for animals treated with N-9 compared to no treatment groups (6-9 days vs 1-9 days for female; 4-6 days vs 1-6 days for males) (Table 1). Finally, we observed more animals still positive at the final day of the experiment (day 20) in the N-9 treated group compared to the no treatment group (100% vs 10% respectively). This was not true for males. We also investigated Luc-Cm DNA shedding in fecal pellets (Supplementary Fig. 2, Supplementary Table 2) and serum antibody elicitation after infection (Supplementary Fig. 3 and 4), showing statistically significant IgA responses in serum after infection regardless of treatment prior to infection. Together these data indicate that treatment with N-9 prior to anorectal infection with Luc-Cm leads to more days of infection and reduces within-group variability for total days and length of infection by IVIS imaging.

**Figure 1.**
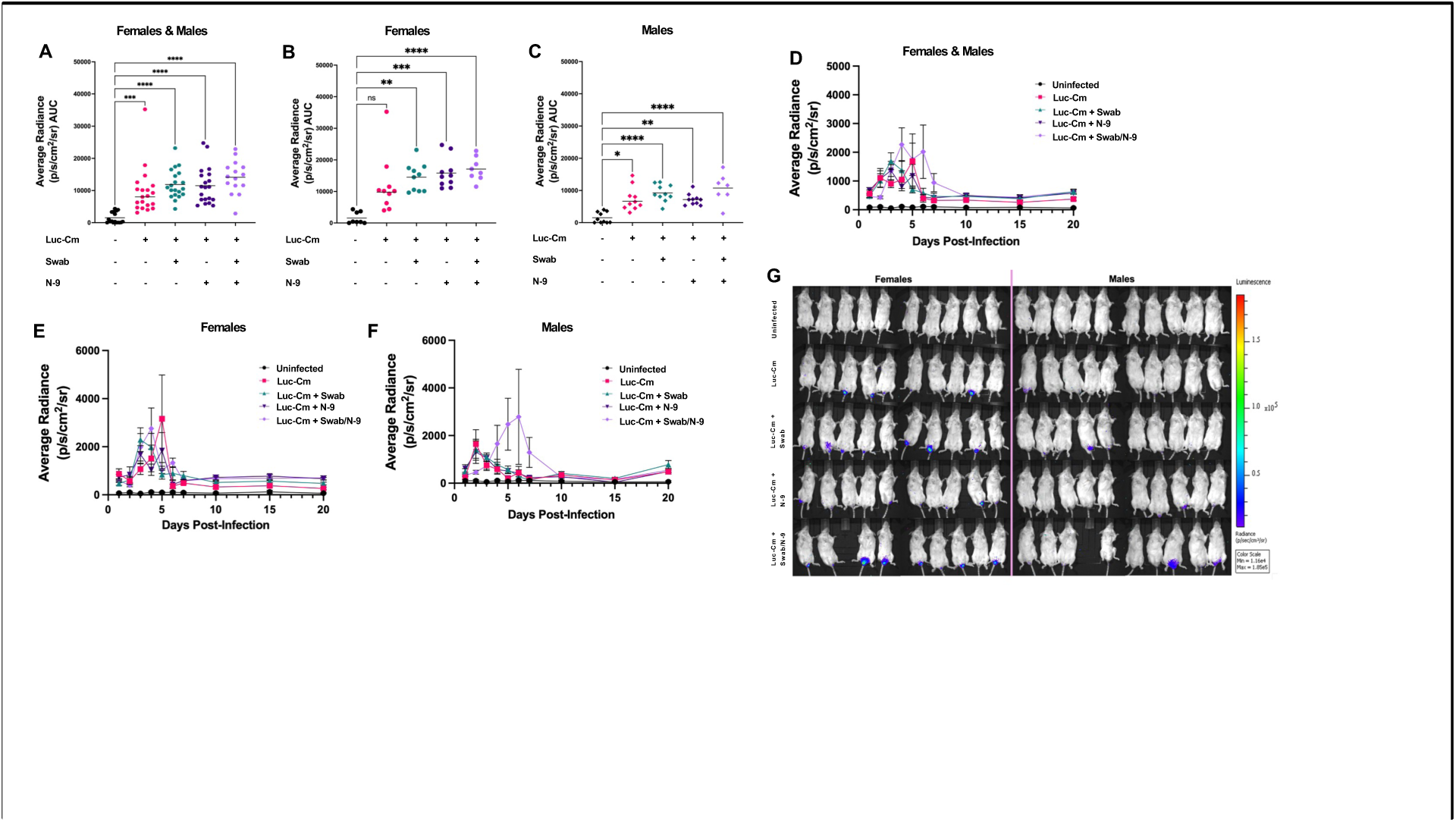
Luc-Cm bacterial burden in the anorectum after epithelial disruption. BALB/c mice (n=20 per group, male and female) were imaged by IVIS after D-luciferin administration anorectally on days 1-20 post infection with Luc-Cm. Average radiance (p/s/cm2/sr) AUC was calculated for each mouse and compared between groups, with either the sexes combined (**A**) or separated into (**B**) females and (**C**) males. The average radiance (p/s/cm2/sr) for each group was throughout the experimental time course post-infection was shown for each group for the (**D**) sexes combined or separated into (**E**) females and (**F**) males. This data was represented as mean ± SEM. (**G**) Representative IVIS images that were taken on day 4 post-infection are shown. Statistical analysis was performed utilizing a non-parametric Kruskal-Wallis One-Way ANOVA. ns = non-significant, * *p* < 0.05, ** *p* < 0.01, *** *p* < 0.001, **** *p* < 0.0001.

**Table 1.**
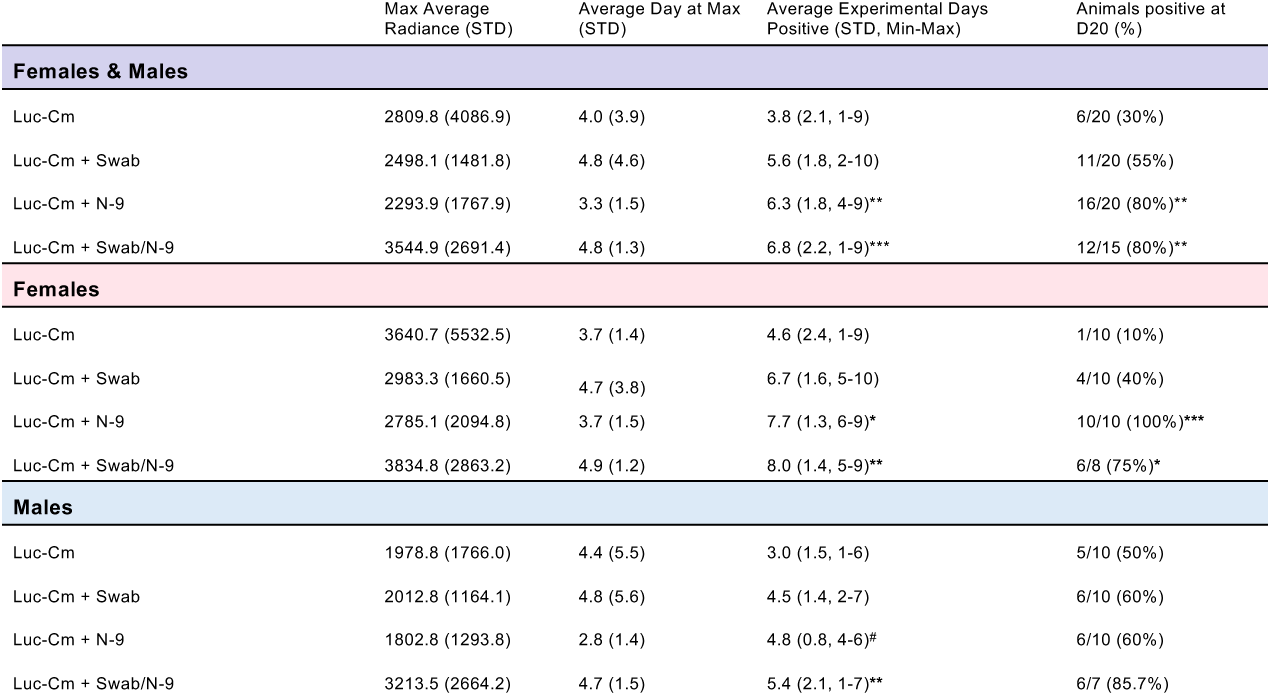
Time course of Luc-Cm bacterial burden. BALB/c mice (n=20 per group, male and female) were imaged by IVIS after D-luciferin administration anorectally on days 1-20 post-infection with Luc-Cm. The average max radiance (p/s/cm2/sr), average day at max radiance, average number of experiment days that were positive (2 times the background radiance), and animals positive at day 20 were described for all of the groups, with either the sexes combined or separated by sex. Statistical analysis was performed utilizing a non-parametric Kruskal-Wallis One-Way ANOVA except for animals positive at day 20, which was analyzed using a Fisher’s exact test. The different epithelial disruption groups were compared to the No Disruption group. * *p* < 0.05, ** *p* < 0.01, *** *p* < 0.001, **** *p* < 0.0001. #The p value was p= 0.0562 between the male No Disruption and Chemical Disruption groups

### 3.2 Combined swab/N-9 treatment prior to anorectal Luc-Cm infection leads to increased morbidity and mortality of mice

Survival, weight, and occurrence of diarrhea were monitored throughout the experiment. A baseline weight for each mouse was recorded on day −4 prior to both epithelial disruption treatment and infection. While several of the mice in the swab/N-9 treatment group displayed notable weight loss (both male and female), none of the treatment groups had statistically significant weight loss compared to the no treatment group (Fig. 2A-C). For females, diarrhea was observed in both the swab (days 0-3, 1 mouse) and swab/N-9 groups (2 mice, day 0 and days 1-3) (Table 2, Fig. 2E). For males, diarrhea was only observed in the swab/N-9 group (5 mice, various days) (Table 2, Fig. 2F). All mice in the no treatment, swab treated, and N-9 treatment groups survived for the duration of the experiment (Fig. 2G-I, Table 2). However, 2 female mice and 3 male mice in the swab/N-9 group died between days 3-15 post-infection (Table 2, Fig. 2G-I). Of these mice, 4 had diarrhea for multiple days prior to death. Additionally, two female mice in the uninfected control group died of unknown causes on day 15 (Fig. 2H). Overall, swab/N-9 combined treatment increased morbidity in infected mice without concomitant increased Luc-Cm infection, suggesting that treatment alone resulted in morbidity.

**Figure 2.**
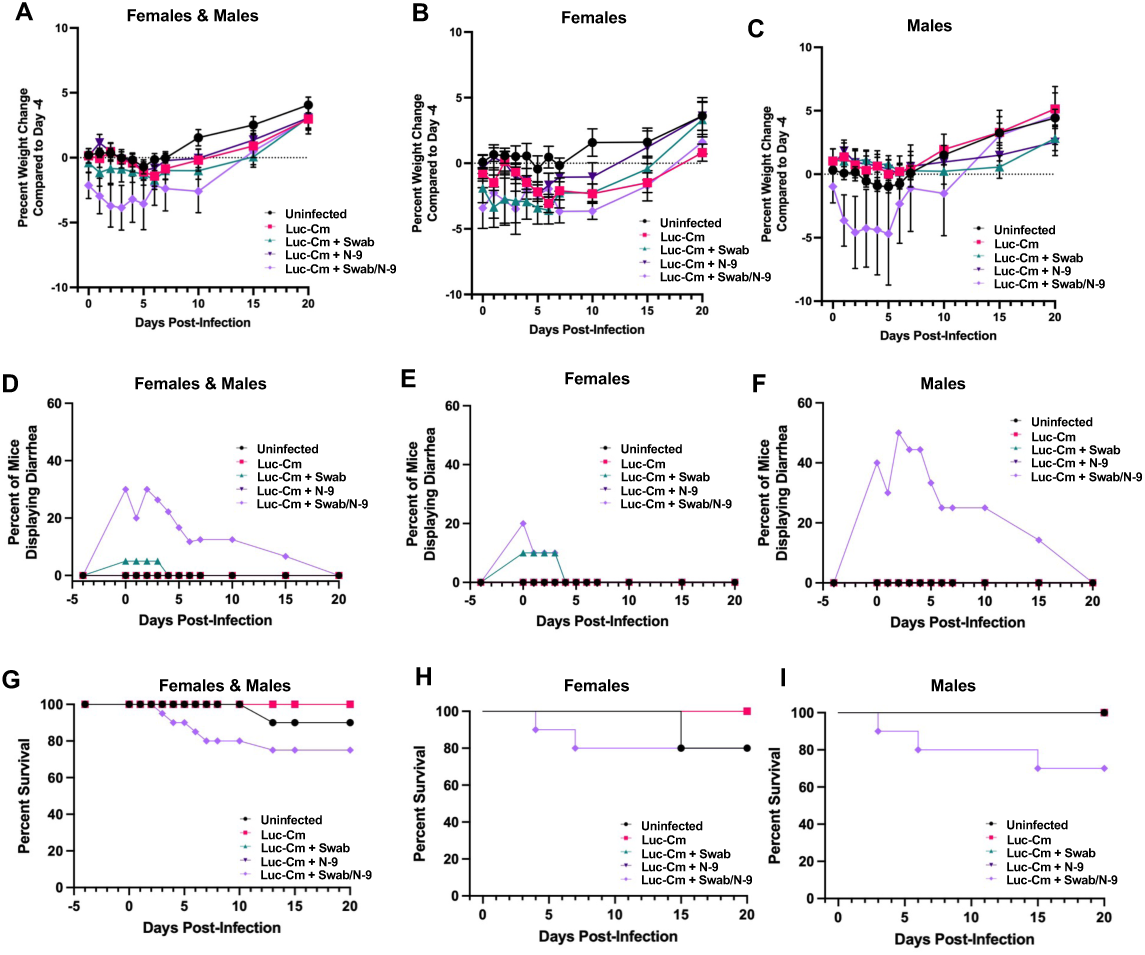
Morbidity and mortality in Luc-Cm anorectally infected mice after prior treatment with epithelial disruption. (**A**) The percent weight change for each mouse was calculated using their initial body weight on day −4 post-infection and means for each group are presented with the SEM error bars for each group. (**B**) The percent of mice in each experimental group with occurrence of diarrhea at each time point are shown. (**C**) The precent survival for each group was monitored throughout the experiment to determine percent survival. Statistical analysis was performed utilizing a Kruskal-Wallis non-parametric one-way ANOVA. ns = non-significant, * *p* < 0.05, ** *p* < 0.01, *** *p* < 0.001, **** *p* < 0.0001.

**Table 2:** Adverse health outcomes after on days 0-20 post-infection. Description of the number of animals that experienced diarrhea and how long that diarrhea lasted for. Also describes the number of animals in the different disruption groups that died during the experimental timeline. The data is display by disruption group, with either the sexes combined or separated. #One of the female mice that died was positive for diarrhea until their final timepoint: All three of the male mice that died were positive for diarrhea until their final timepoint.

|  | Maximum number of<br>animals positive for diarrhea | Range of days<br>positive for diarrhea | Number of<br>animals that died |
| --- | --- | --- | --- |
| **Females & Males** |  |  |  |
| Luc-Cm | 0 | N/A | 0 |
| Luc-Cm + Swab | 1 | 4 days | 0 |
| Luc-Cm + N-9 | 0 | N/A | 0 |
| Luc-Cm + Swab/N-9 | 6 | 1 to 16 days | 5<sup>#</sup> |
| **Females** |  |  |  |
| Luc-Cm | 0 | N/A | 0 |
| Luc-Cm + Swab | 1 | Days 0-3 | 0 |
| Luc-Cm + N-9 | 0 | N/A | 0 |
| Luc-Cm + Swab/N-9 | 2 | Days 0-3 | 2<sup>#</sup> |
| **Males** |  |  |  |
| Luc-Cm | 0 | N/A | 0 |
| Luc-Cm + Swab | 0 | N/A | 0 |
| Luc-Cm + N-9 | 0 | N/A | 0 |
| Luc-Cm + Swab/N-9 | 5 | Days 1-15 | 3<sup>#</sup> |

### 3.3 Epithelial changes after N-9 treatment

N-9 treatment prior to inoculation of Luc-Cm in the anorectal tract of mice showed decreased within-group variability of infection as detected by IVIS. To further explore the impact of N-9 treatment on the anorectal epithelium, we examined epithelial changes after N-9 treatment in the distal colon using histological examination and examination of fecal pellets in uninfected animals (Fig. 3). 24 hours after treatment with N-9, mouse fecal pellets were collected and the distal colon was removed and fixed for histological staining. Mucin-2 (MUC2), a marker of goblet cells, was decreased in N-9 treated animals compared to untreated animals, suggesting that N-9 treatment led to loss of goblet cells (Fig. 3A-C). Fecal pellets from N-9 treated animals showed increased presence of IgA (Fig. 3D-F) and Lipocalin-2 (LCN-2) (Fig. 3G-I), indicators of epithelial barrier disruption and intestinal inflammation respectively. Together, these results indicate that N-9 treatment may allow for more robust Luc-Cm infection of epithelial cells by disrupting the mucosal barrier and intestinal integrity in the anorectal tract.

**Figure 3:**
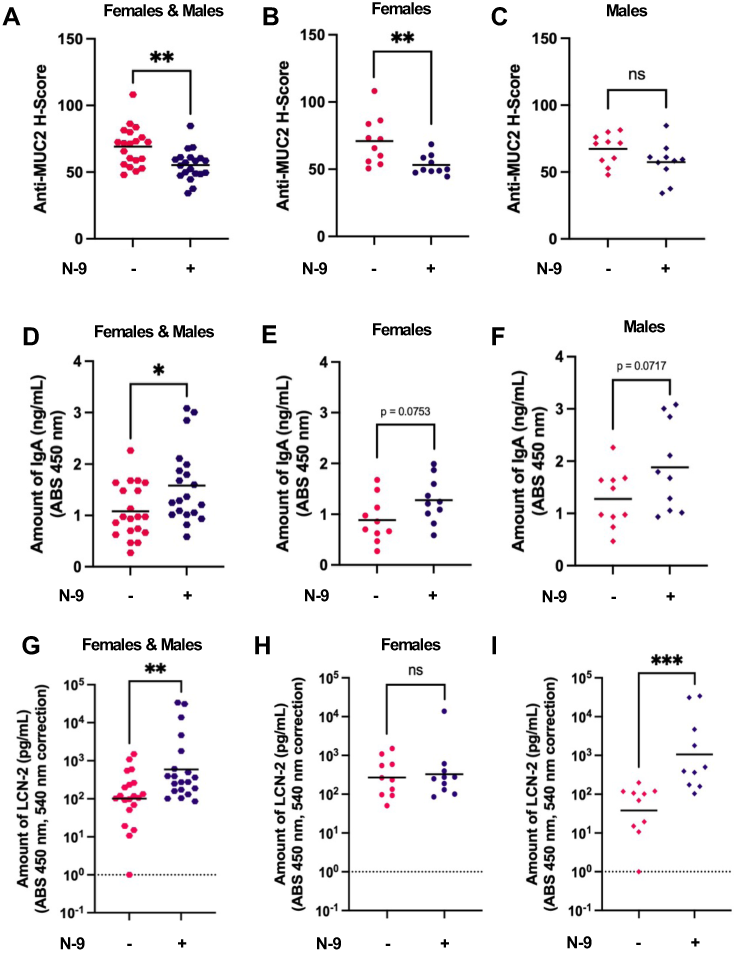
BALB/c mice (n=20) were mock vaccinated with PBS 3-times, 3-weeks apart. On day 68, the mice received either no disruption (n=20) or chemical disruption (n=20) treatment in the anorectal tract. The next day fecal material was collected, the mice were euthanized, and the distal colon was collected for histology. For IHC, the H-score of distal colons that were stained with an anti-MUC2 antibody were reported for each group with either the sexes combined (**A**) or separated in females (**B**) and males (**C**). H-score accounts for the number of cells that stain positive for the antibody and how positively the cells are stained. The amount of IgA (ng/mL) in fecal material was determined using an ELISA kit for each group with either the sexes combined (**D**) or separated in females (**E**) and males (**F**). The amount of LCN-2 (pg/mL) in fecal material was determined using an ELISA kit for each group with either the sexes combined (**G**) or separated in females (**H**) and males (**I**). Statistical analysis was performed utilizing a non-parametric Mann-Whitney t-test. ns = non-significant, * *p* < 0.05, ** *p* < 0.01, *** *p* < 0.001, **** *p* < 0.0001.

### 3.4 Examining vaccine efficacy against anorectal Luc-Cm infection

Having identified that N-9 treatment prior to anorectal inoculation with Luc-Cm resulted in more days of infection and reduced within-group variability for total days of and length of infection using IVIS imaging without concomitant increase in morbidity, we next used this model to investigate the efficacy of a vaccine candidate our lab has shown to provide protection in a mouse FRT *Chlamydia* infection model (data not shown). This vaccine is composed of a short peptide corresponding to a highly conserved epitope in the variable domain 4 (VD4) of the *Chlamydia muridarum* major outer membrane protein (MOMP)[1]. Male and female BALB/c mice (n = 20/group) were immunized intramuscularly 3 times, 3-weeks apart, with either Qβ-VD4-Cm or Qβ control. An additional group of mice were included as an uninfected control. Sera and fecal pellets were collected prior to infection to monitor vaccine-elicited antibody titers against the VD4-Cm peptide. Immunization with Qβ-VD4-Cm elicited consistently high-titer serum IgG against the VD4-Cm peptide (Fig. 4A-C). In addition, serum IgA (Fig. 4D-F) and fecal IgG (Fig. 4G-I) against the VD4-Cm peptide were detected in several mice. However, there was no significant induction of fecal IgA against the VD4-Cm peptide in Qβ-VD4-Cm immunized animals (Fig. 4J-L).

**Figure 4:**
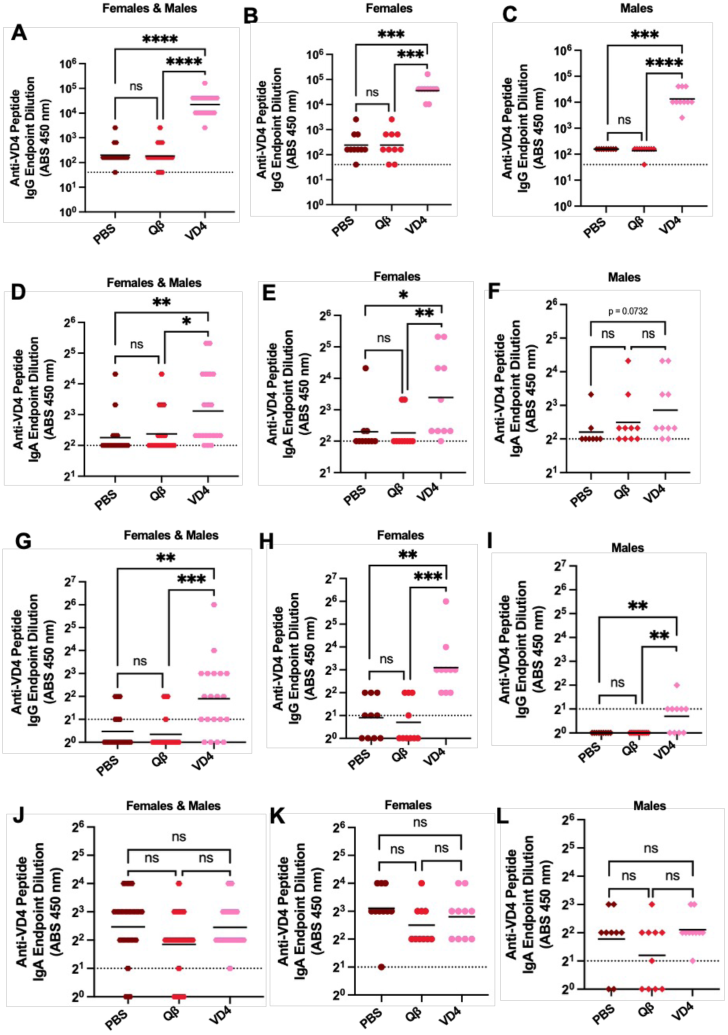
Immune sera and fecal pellets from immunized mice were collected at 3-weeks post-3^rd^ immunization and the antibody binding capacity to the Cm VD4 peptide was determined using ELISA. (**C**) The sera IgG endpoint dilution titers were also shown for each group, with either the (**A**) sexes combined or separated into (**B**) females or males. The serum IgA bind capacity were shown as endpoint dilution titers for each group, with either the (**D**) sexes combined or separated into (**E**) females or (**F**) males. The fecal IgG bind capacity were shown as endpoint dilution titers for each group, with either the (**G**) sexes combined or separated into (**H**) females or (**I**) males. The fecal IgA bind capacity was shown as endpoint dilution titers for each group, with either the (**J**) sexes combined or separated into (**K**) females or (**L**) males. Statistical analysis was performed utilizing a Kruskal-Wallis non-parametric one-way ANOVA. ns = non-significant, * *p* < 0.05, ** *p* < 0.01, *** *p* < 0.001, **** *p* < 0.0001.

Both the Qβ-VD4-Cm and Qβ immunized mice had similar infection time course, with the infection peaking on day 3-4 and then tapering off for the remainder of the experimental days (Table 3, Fig. 5D-F). Immunization with Qβ-VD4-Cm did not decrease Luc-Cm bacterial burden compared to the Qβ control as measured by IVIS (Fig. 5A-C, Fig. 5G), despite the production of high-titer, VD4-Cm serum IgG antibodies (Fig. 4A-C) These results show that our Qβ-VD4-Cm vaccine was unable to decrease Luc-Cm bacterial burden in the anorectal tract, despite previously being shown to reduce infection at early time points in the FRT.

**Figure 5:**
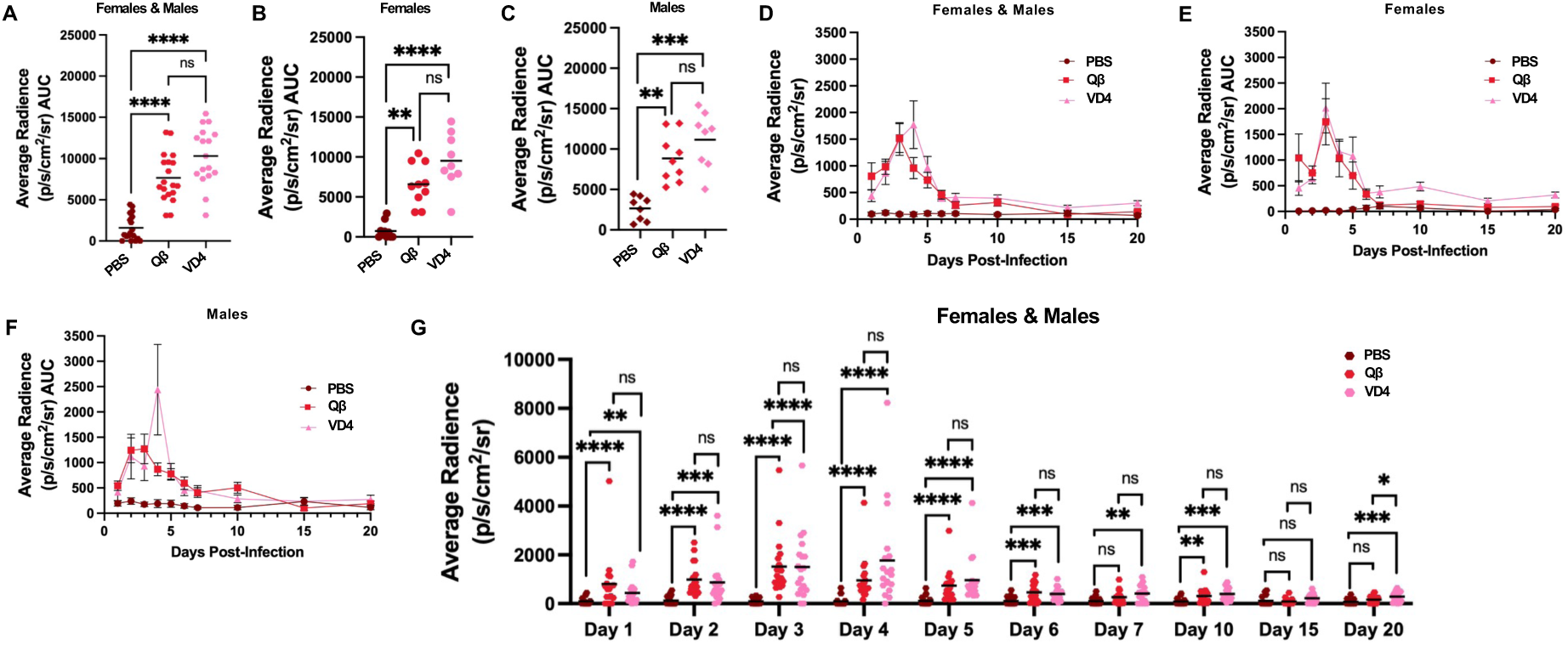
Luc-Cm bacterial burden in the anorectum after chemical disruption of immunized mice. Immunized BALB/c mice (n=20 per group, male and female) were imaged by IVIS after D-luciferin administration anorectally on days 1-20 post-infection with Luc-Cm. Average radiance (p/s/cm2/sr) AUC for days 1 to 20 post-infection was calculated for each mouse and compared between the different immunization groups, with either the (**A**) sexes combined or separated into (**B**) females or (**C**) males.. Average radiance (p/s/cm2/sr) time course for days 1 to 20 post-infection was shown for the different immunization groups, with either the (**D**) sexes combined or separated into (**E**) females or (**F**) males. (**G**) The average radiance (p/s/cm2/sr) for each mouse in the different immunization groups was shown for days 1 to 20 post-infection for both of the sexes combined. Statistical analysis was performed utilizing a non-parametric Kruskal-Wallis One-Way ANOVA. ns = non-significant, * *p* < 0.05, ** *p* < 0.01, *** *p* < 0.001, **** *p* < 0.0001.

**Table 3.** Time course of Luc-Cm bacterial burden in immunized mice after chemical disruption. BALB/c mice (n=20 per group, male and female) were imaged by IVIS after D-luciferin administration anorectally on days 1-20 post-infection with Luc-Cm. The average max radiance (p/s/cm2/sr), average day at max radiance, average number of experiment days that were positive (2 times the background radiance), and animals positive at day 20 were described for all of the groups, with either the sexes combined or separated by sex. Statistical analysis was performed utilizing a non-parametric Mann-Whitney t-test except for animals positive at day 20, which was analyzed using a Fisher’s exact test. The different epithelial disruption groups were compared to the QB group. * *p* < 0.05, ** *p* < 0.01, *** *p* < 0.001, **** *p* < 0.0001.

|  | Max Average<br>Radiance (STD) | Average Day at<br>Max (STD) | Average experimental<br>days positive (STD, Min-Max) | Animals positive<br>at D20 (%) |
| --- | --- | --- | --- | --- |
| **Females & Males** |  |  |  |  |
| Qβ Control | 2148.5 (1423.8) | 3.2 (1.4) | 4.5 (1.5, 1-7) | 0/19 (0%) |
| Qβ-VD4-Cm | 2892.9 (1884.5) | 3.6 (0.9) | 4.4 (1.5, 2-7) | 2/17 (11.8%) |
| **Females** |  |  |  |  |
| Qβ Control | 2577.5 (1739.6) | 2.9 (1.3) | 3.6 (1.0, 2-5) | 0/10 (0%) |
| Qβ-VD4-Cm | 2554.4 (1444.6) | 3.7 (0.9) | 4.3 (1.4, 2-6) | 1/10 (10%) |
| **Males** |  |  |  |  |
| Qβ Control | 1672.0 (819.4) | 3.6 (1.5) | 5.4 (1.3, 3-7) | 0/7 (0%) |
| Qβ-VD4-Cm | 3376.4 (2424.1) | 3.4 (0.8) | 4.6 (1.6, 3-7) | 1/7 (14.3%) |

## 4. Discussion

The anorectal tract is an important anatomical site of Ct infection. Relevant anorectal infection models are needed to test the efficacy of vaccine candidates and antibiotic treatments at the anorectal site. This is crucial as treatment efficacy can differ between anatomical sites, which is demonstrated by the difference in effectiveness of the antibiotic azithromycin at the urogenital and anorectal tract [4,28]. Here, we examined how disrupting the anorectal epithelium prior to anorectal inoculation with Luc-Cm impacts the progression of infection. We found that N-9 treatment was able to decrease the within-group variability of infection without concomitant diarrhea, weight loss, or mortality. Additionally, we assessed the efficacy of intramuscular immunization with Qβ-VD4-Cm in this model, finding that immunization Qβ-VD4-Cm was insufficient to decrease Luc-Cm infection in the anorectal tract despite eliciting high-titer IgG in the sera.

N-9 is a spermicide that is commercially available in different birth control products. Pretreatment with N-9 has been associated with an increased risk of both HPV and HIV in humans [29]. Indeed, N-9 pretreatment is a common strategy to increase viral load and reproducibility of HPV infections in mouse infection models [17,18]. N-9 treatment has been used to facilitate HPV infection in both the vaginal and anorectal tract of mice [17,18]. This is potentially due to N-9 causing epithelial disruption at these sites. For HPV, it is hypothesized that epithelial disruption due to pretreatment with N-9 mimics the epithelial micro-abrasions that occur during sexual activity, which allows HPV access to basal epithelial cells for infection [26,27]. However, it is unclear exactly how N-9 is promoting Cm infection consistency in the anorectal tract. As Cm can infect mucosal epithelial cells closer to the surface, infection is unlikely to require micro-abrasions to access suitable host cells. However, there are several potential ways that N-9 could be improving Cm infection. N-9 could disrupt the mucus layer to increase Cm access to epithelial cells [28]. Disruption of the mucosa could also limit mucus trapping, and subsequent clearance, of Cm during inoculation [29]. N-9 could also be directly causing epithelial cell sloughing and decreasing intestinal barrier function [27]. This could promote an immune response that is skewed towards tissue repair instead of anti-microbial functions at the 24-hour time point that the mice were examined. In our model, we found that N-9 treatment in mice resulted in decreased goblet cells (Fig. 3), indicating potential disruption of the mucosal barrier, and increased markers of intestinal barrier disruption. However, additional work is needed to fully characterize the immune environment and changes to the mucosal composition of the anorectal tract after N-9 treatment.

Throughout the experiments, we examined for potential sex difference within the treatment groups and observed some significant differences in bacterial burden and shedding in the model establishment experiments when comparing male and female mice. These differences were not surprising as the gastrointestinal microbiome of female and male mice starts diverging after sexual maturity, which could impact infection [30–32]. Overall, female mice tended to have higher bacterial burden and shedding when compared to males in the same group. The higher burden in female mice might be due to the potential for cross-over infections back and forth with the FRT [30–32]. However, we did not see a significant increased bacterial burden in female mice immunized with Qβ Control or Qβ-VD4-Cm, compared to male mice. To try to minimize the impact of potential crossover infections with the urogenital tract on the average radiance detected by IVIS, D-luciferin was directly administered into the anorectal tract 3 minutes prior to imaging [4,8,35]. While D-luciferin is typically administered using intraperitonially (I.P.) injection, unpublished work from our lab has indicated that similar luminescence is detected if D-luciferin is instilled directly in the rectum (data not shown) [36]. An additional sex difference we noticed was female mice immunized with Qβ-VD4-Cm had higher vaccine-elicited IgG antibody titers in sera and fecal material, compared to male mice immunized with Qβ-VD4-Cm. This was anticipated as females tend to have a stronger antibody response compared to males in response to both natural infections and vaccination [32].

Currently, there are limited animal models of *Chlamydia* infections that do not focus on ocular or FRT as the site of infection. As *Chlamydia* can infect various anatomical locations, it is important to develop infection models for these sites of infection to evaluate potential differences in immune response and vaccine efficacy [15,34]. Our work here could provide an additional basis to support the development of mouse models of *Chlamydia* infection at various anatomical sites.

We did not see vaccine-elicited protection in our model of anorectal tract infection, despite our Qβ-VD4-Cm vaccine providing protection in the vaginal tract of BALB/c mouse. This could be due to a several factors. Intramuscular immunization typically promotes a strong systemic IgG response; however, higher vaccine-elicited IgA titers might promote increased efficacy in the anorectal tract [21,36]. Future work can be done to increase vaccine-elicited IgA titers to provide protection at both the vaginal tract and the anorectal tract, such as utilizing vaccine adjuvants.

Changing the route of immunization can also alter the antibody subtypes produced to improve protection at mucosal sites [21,36]. Intranasal immunization of bacteriophage VLP vaccines has been shown to increase sera IgA titers [21]. Also, N-9 treatment in the anorectal tract seems to be disrupting the mucosal barrier in our model, disruptions to the anal mucosa could be limiting mucosal trapping even in the presence of vaccine-elicited antibodies [29].

In conclusion, our results show that N-9 pretreatment of the anorectum can decrease the within-group variation and improve persistence of Luc-Cm infection in mice. Our results highlight the importance of examining *Chlamydia* vaccine efficacy at multiple relevant sites as there may be anatomical limitations of protection.

## 5. Acknowledgements

We would like to thank Dr. Guangming Zhong for sharing the luciferase-expressing *Chlamydia muridarum* (Luc-Cm).

## References

1. Collar, A.L.; Linville, A.C.; Core, S.B.; Frietze, K.M. Epitope-Based Vaccines against the Chlamydia Trachomatis Major Outer Membrane Protein Variable Domain 4 Elicit Protection in Mice. Vaccines 2022, 10, 875, doi:10.3390/vaccines10060875.

2. Sinka, K. The Global Burden of Sexually Transmitted Infections. Clin. Dermatol. 2024, 42, 110–118, doi:10.1016/j.clindermatol.2023.12.002.

3. Yonke, N.; Aragón, M.; Phillips, J.K. Chlamydial and Gonococcal Infections: Screening, Diagnosis, and Treatment. 2022, 105.

4. Khosropour, C.M.; Dombrowski, J.C.; Vojtech, L.; Patton, D.L.; Campbell, L.A.; Barbee, L.A.; Franzi, M.C.; Hybiske, K. Rectal Chlamydia Trachomatis Infection: A Narrative Review of the State of the Science and Research Priorities. Sex. Transm. Dis. 2021, 48, e223–e227, doi:10.1097/OLQ.0000000000001549.

5. Ceccarani, C.; Gaspari, V.; Morselli, S.; Djusse, M.E.; Venturoli, S.; Camboni, T.; Severgnini, M.; Foschi, C.; Consolandi, C.; Marangoni, A. Chlamydia Trachomatis and Neisseria Gonorrhoeae Rectal Infections: Interplay between Rectal Microbiome, HPV Infection and Torquetenovirus. PLOS ONE 2024, 19, e0301873, doi:10.1371/journal.pone.0301873.

6. Assi, R. Sexually Transmitted Infections of the Anus and Rectum. World J. Gastroenterol. 2014, 20, 15262, doi:10.3748/wjg.v20.i41.15262.

7. Smolarczyk, K.; Młynarczyk-Bonikowska, B.; Majewski, S.; Rudnicka, E.; Unemo, M.; Fiedor, P. Lymphogranuloma Venereum: An Emerging Problem in Poland. Adv. Dermatol. Allergol. 2022, 39, 454–459, doi:10.5114/ada.2020.101466.

8. Wang, L.; Zhang, Q.; Zhang, T.; Zhang, Y.; Zhu, C.; Sun, X.; Zhang, N.; Xue, M.; Zhong, G. The Chlamydia Muridarum Organisms Fail to Auto-Inoculate the Mouse Genital Tract after Colonization in the Gastrointestinal Tract for 70 Days. PLOS ONE 2016, 11, e0155880, doi:10.1371/journal.pone.0155880.

9. Yeruva, L.; Spencer, N.; Bowlin, A.K.; Wang, Y.; Rank, R.G. Chlamydial Infection of the Gastrointestinal Tract: A Reservoir for Persistent Infection. Pathog. Dis. 2013, 68, 88–95, doi:10.1111/2049-632X.12052.

10. Ma, J.; Zhang, T.; Tian, Q.; Xiao, M.; Han, L.; Liu, Q.; Zhong, G.; Liu, Y. Chlamydia Muridarum Mutant CM-pGP3S as a Novel Attenuated Live Rectal Vaccine Protects against Genital Tract Infection. Front. Cell. Infect. Microbiol. 2025, 15, 1550455, doi:10.3389/fcimb.2025.1550455.

11. Mishkin, N.; Carrasco, S.E.; Palillo, M.; Momtsios, P.; Woods, C.; Henderson, K.S.; Longhini, A.L.F.; Otis, C.; Gardner, R.; Joseph, A.M.;, et al. Chlamydia Muridarum Causes Persistent Subclinical Infection and Elicits Innate and Adaptive Immune Responses in C57BL/6J, BALB/cJ, and J:ARC(S) Mice Following Exposure to Shedding Mice. Comp. Med. 2024, 74, 373–391, doi:10.30802/AALAS-CM-24-057.

12. Mishkin, N.; Ricart Arbona, R.J.; Carrasco, S.E.; Lawton, S.; Henderson, K.S.; Momtsios, P.; Sigar, I.M.; Ramsey, K.H.; Cheleuitte-Nieves, C.; Monette, S.;, et al. Reemergence of the Murine Bacterial Pathogen *Chlamydia Muridarum* in Research Mouse Colonies. Comp. Med. 2022, 72, 230–242, doi:10.30802/AALAS-CM-22-000045.

13. Palillo, M.B.; Mishkin, N.; Aydin, M.; Mourino, A.; Ricart Arbona, R.J.; Lipman, N.S. Examining Intercage Transmission of Chlamydia Muridarum: Impact of Barrier Husbandry and Cage Sanitization. J. Am. Assoc. Lab. Anim. Sci. 2024, 63, 513–520, doi:10.30802/AALAS-JAALAS-24-043.

14. Campbell, J.; Huang, Y.; Liu, Y.; Schenken, R.; Arulanandam, B.; Zhong, G. Bioluminescence Imaging of Chlamydia Muridarum Ascending Infection in Mice. PLoS ONE 2014, 9, e101634, doi:10.1371/journal.pone.0101634.

15. Brunham, R.C.; Rey-Ladino, J. Immunology of Chlamydia Infection: Implications for a Chlamydia Trachomatis Vaccine. Nat. Rev. Immunol. 2005, 5, 149–161, doi:10.1038/nri1551.

16. Slade, J.A.; Hall, J.V.; Kintner, J.; Phillips-Campbell, R.; Schoborg, R.V. Host Nectin-1 Promotes Chlamydial Infection in the Female Mouse Genital Tract, but Is Not Required for Infection in a Novel Male Murine Rectal Infection Model. PLOS ONE 2016, 11, e0160511, doi:10.1371/journal.pone.0160511.

17. Cladel, N.M.; Budgeon, L.R.; Balogh, K.K.; Cooper, T.K.; Hu, J.; Christensen, N.D. A Novel Pre-Clinical Murine Model to Study the Life Cycle and Progression of Cervical and Anal Papillomavirus Infections. PLOS ONE 2015, 10, e0120128, doi:10.1371/journal.pone.0120128.

18. Roberts, J.N.; Buck, C.B.; Thompson, C.D.; Kines, R.; Bernardo, M.; Choyke, P.L.; Lowy, D.R.; Schiller, J.T. Genital Transmission of HPV in a Mouse Model Is Potentiated by Nonoxynol-9 and Inhibited by Carrageenan. Nat. Med. 2007, 13, 857–861, doi:10.1038/nm1598.

19. Andrew, D.W.; Cochrane, M.; Schripsema, J.H.; Ramsey, K.H.; Dando, S.J.; O’Meara, C.P.; Timms, P.; Beagley, K.W. The Duration of Chlamydia Muridarum Genital Tract Infection and Associated Chronic Pathological Changes Are Reduced in IL-17 Knockout Mice but Protection Is Not Increased Further by Immunization. PLoS ONE 2013, 8, e76664, doi:10.1371/journal.pone.0076664.

20. Webster, E.; Seiger, K.W.; Core, S.B.; Collar, A.L.; Knapp-Broas, H.; Graham, J.; Shrestha, M.; Afzaal, S.; Geisler, W.M.; Wheeler, C.M.;, et al. Immunogenicity and Protective Capacity of a Virus-like Particle Vaccine against Chlamydia Trachomatis Type 3 Secretion System Tip Protein, CT584. Vaccines 2022, 10, 111, doi:10.3390/vaccines10010111.

21. Jamus, A.N.; Wilton, Z.E.R.; Armijo, S.D.; Flanagan, J.; Romano, I.G.; Core, S.B.; Frietze, K.M. Nasal and Ocular Immunization with Bacteriophage Virus-like Particle Vaccines Elicits Distinct Systemic and Mucosal Antibody Profiles. Vaccines 2025, 13, 829, doi:10.3390/vaccines13080829.

22. Gerber, P.F.; Opriessnig, T. Detection of Immunoglobulin (Ig) A Antibodies against Porcine Epidemic Diarrhea Virus (PEDV) in Fecal and Serum Samples. MethodsX 2015, 2, 368–373, doi:10.1016/j.mex.2015.10.001.

23. Erben, U.; Loddenkemper, C.; Doerfel, K.; Spieckermann, S.; Haller, D.; Heimesaat, M.; Zeitz, M.; Siegmund, B.; Kühl, A.A. A Guide to Histomorphological Evaluation of Intestinal Inflammation in Mouse Models.

24. Dombrowski, J.C.; Wierzbicki, M.R.; Newman, L.M.; Powell, J.A.; Miller, A.; Dithmer, D.; Soge, O.O.; Mayer, K.H. Doxycycline Versus Azithromycin for the Treatment of Rectal Chlamydia in Men Who Have Sex With Men: A Randomized Controlled Trial. Clin. Infect. Dis. 2021, 73, 824–831, doi:10.1093/cid/ciab153.

25. Phillips, D.M.; Sudol, K.M.; Taylor, C.L.; Guichard, L.; Elsen, R.; Maguire, R.A. Lubricants Containing N-9 May Enhance Rectal Transmission of HIV and Other STIs. Contraception 2004, 70, 107–110, doi:10.1016/j.contraception.2004.04.008.

26. Frietze, K.M.; Lijek, R.; Chackerian, B. Applying Lessons from Human Papillomavirus Vaccines to the Development of Vaccines against *Chlamydia Trachomatis*. Expert Rev. Vaccines 2018, 17, 959–966, doi:10.1080/14760584.2018.1534587.

27. Fuchs, E.J.; Grohskopf, L.A.; Lee, L.A.; Bakshi, R.P.; Hendrix, C.W. Quantitative Assessment of Altered Rectal Mucosal Permeability Due to Rectally Applied Nonoxynol-9, Biopsy, and Simulated Intercourse. J. Infect. Dis. 2013, 207, 1389–1396, doi:10.1093/infdis/jit030.

28. Sheng, Y.H.; Hasnain, S.Z. Mucus and Mucins: The Underappreciated Host Defence System. Front. Cell. Infect. Microbiol. 2022, 12, 856962, doi:10.3389/fcimb.2022.856962.

29. Schaefer, A.; Lai, S.K. The Biophysical Principles Underpinning Muco-Trapping Functions of Antibodies. Hum. Vaccines Immunother. 2022, 18, 1939605, doi:10.1080/21645515.2021.1939605.

30. McGee, J.S.; Huttenhower, C. Of Mice and Men and Women: Sexual Dimorphism of the Gut Microbiome. Int. J. Womens Dermatol. 2021, 7, 533–538, doi:10.1016/j.ijwd.2021.10.007.

31. Yurkovetskiy, L.; Burrows, M.; Khan, A.A.; Graham, L.; Volchkov, P.; Becker, L.; Antonopoulos, D.; Umesaki, Y.; Chervonsky, A.V. Gender Bias in Autoimmunity Is Influenced by Microbiota. Immunity 2013, 39, 400–412, doi:10.1016/j.immuni.2013.08.013.

32. Fischinger, S.; Boudreau, C.M.; Butler, A.L.; Streeck, H.; Alter, G. Sex Differences in Vaccine-Induced Humoral Immunity. Semin. Immunopathol. 2019, 41, 239–249, doi:10.1007/s00281-018-0726-5.

33. Khalil, A.A.; Jameson, M.J.; Broaddus, W.C.; Chung, T.D.; Golding, S.E.; Dever, S.M.; Rosenberg, E.; Valerie, K. Subcutaneous Administration of D-Luciferin Is an Effective Alternative to Intraperitoneal Injection in Bioluminescence Imaging of Xenograft Tumors in Nude Mice. ISRN Mol. Imaging 2013, 2013, 1–7, doi:10.1155/2013/689279.

34. Elwell, C.; Mirrashidi, K.; Engel, J. Chlamydia Cell Biology and Pathogenesis. Nat. Rev. Microbiol. 2016, 14, 385–400, doi:10.1038/nrmicro.2016.30.

35. Bryan, E.R.; Redgrove, K.A.; Mooney, A.R.; Mihalas, B.P.; Sutherland, J.M.; Carey, A.J.; Armitage, C.W.; Trim, L.K.; Kollipara, A.; Mulvey, P.B.M.;, et al. Chronic Testicular Chlamydia Muridarum Infection Impairs Mouse Fertility and Offspring Development†. Biol. Reprod. 2020, 102, 888–901, doi:10.1093/biolre/ioz229.

36. Pabst, O.; Slack, E. IgA and the Intestinal Microbiota: The Importance of Being Specific. Mucosal Immunol. 2020, 13, 12–21, doi:10.1038/s41385-019-0227-4.

37. Sha, H.; He, X.; Yan, K.; Li, J.; Li, X.; Xie, Y.; Yang, Y.; Deng, Y.; Li, G.; Yang, J. Blocking Coprophagy Increases the Levels of Inflammation and Depression in Healthy Mice as Well as Mice Receiving Fecal Microbiota Transplantation from Disease Model Mice Donors. APMIS 2023, 131, 351–368, doi:10.1111/apm.13326.

