## Supplementary figures for "Assessment of *in vivo* epithelial disruption to model anorectal *Chlamydia* infection in mice"

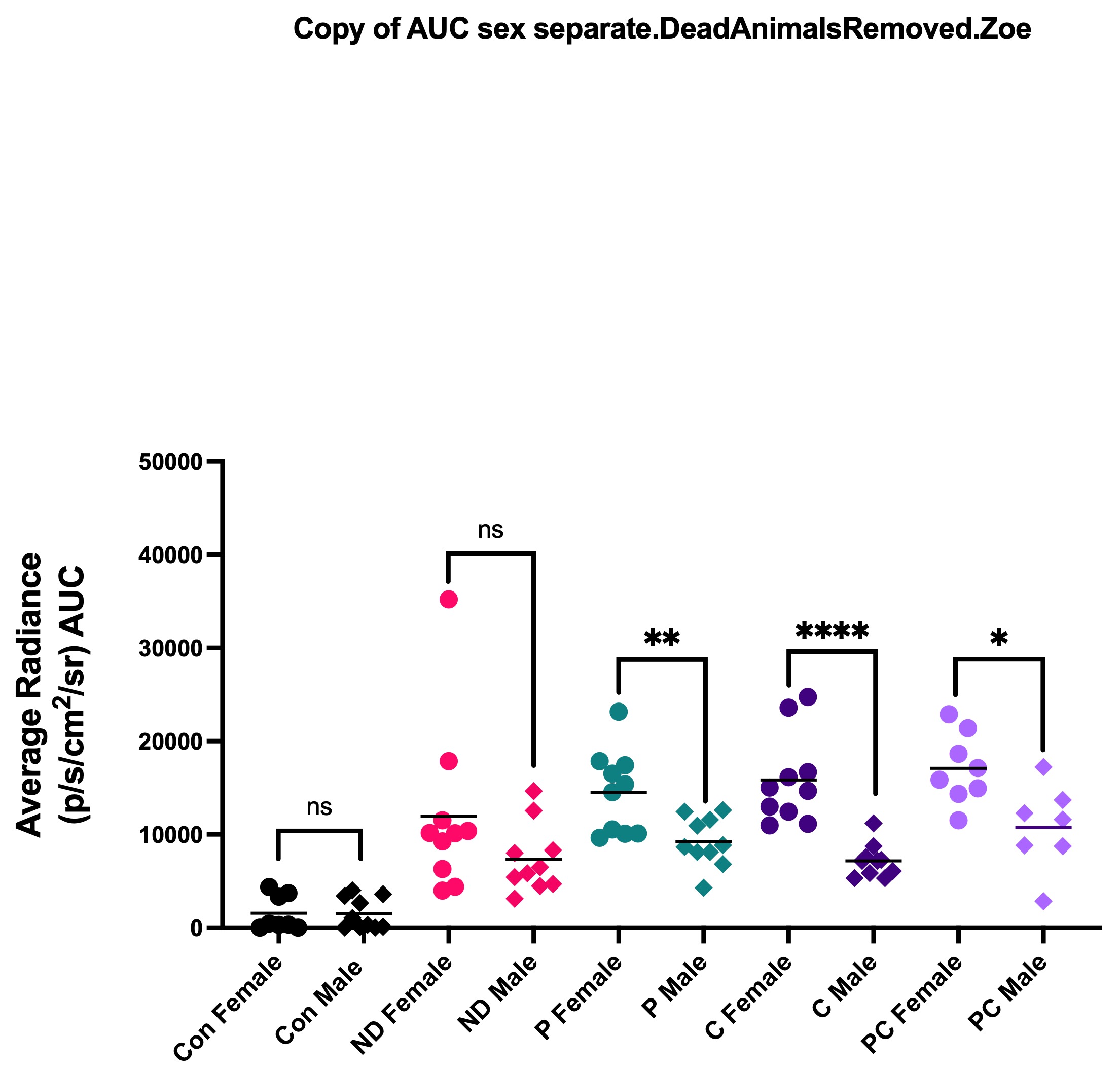

**Females vs. Males**

**Uninfected (F)**

**Uninfected (M)**

**Luc-Cm (F)**

**Luc-Cm (M)**

**Luc-Cm + Swab (F)**

**Luc-Cm + Swab (M)**

**Luc-Cm + N-9 (F)**

**Luc-Cm + N-9 (M)**

**Luc-Cm + Swab/N-9 (M)**

**Luc-Cm + Swab/N-9 (F)**

**Supplementary Figure 1**: **Sex differences in Luc-Cm bacterial burden in the anorectum after epithelial disruption.** BALB/c mice (n=20 per group, male and female) were imaged by IVIS after D-luciferin administration anorectally on days 1-20 post-infection with Luc-Cm. Average radiance (p/s/cm2/sr) AUC was calculated for each mouse and compared between females and males from the same disruption groups. Statistical analysis was performed utilizing a non-parametric Mann-Whitney t-test. ns = non-significant, * *p* < 0.05, ** *p* < 0.01, *** *p* < 0.001, **** *p* < 0.0001

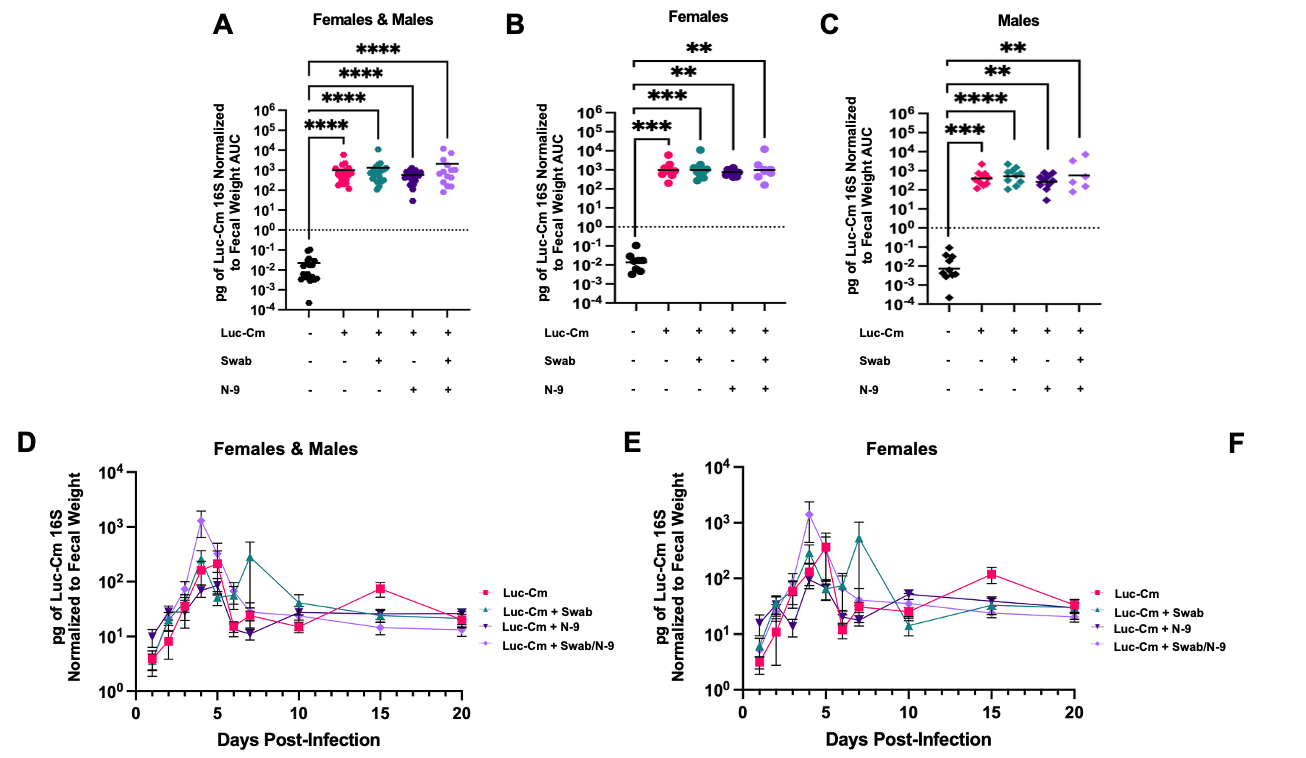

**Supplementary Figure 2. Luc-Cm shedding in feces after epithelial disruption**. Feces from BALB/c mice (n=20/group, male and female) were collected on days 0-20 after anorectal infection with Luc-Cm. Feces were process to extract DNA and RealTime-qPCR was used to determine the pg of Luc-Cm 16S normalized to fecal weight using the formula (C × VT)/(VU × M). AUC was calculated for each mouse and compared between groups, with either the (**A**) sexes combined or separated into (**B**) females or (**C**) males. The average pg of Luc-Cm 16S normalized to fecal weight was shown for each group throughout the experimental time course post-infection, with either the (**D**) sexes combined or separated into (**E**) females or (**F**) males. This data was represented as mean ± SEM. Statistical analysis was performed utilizing a non-parametric Kruskal-Wallis One-Way ANOVA. ns = non-significant, * *p* < 0.05, ** *p* < 0.01, *** *p* < 0.001, **** *p* < 0.0001.

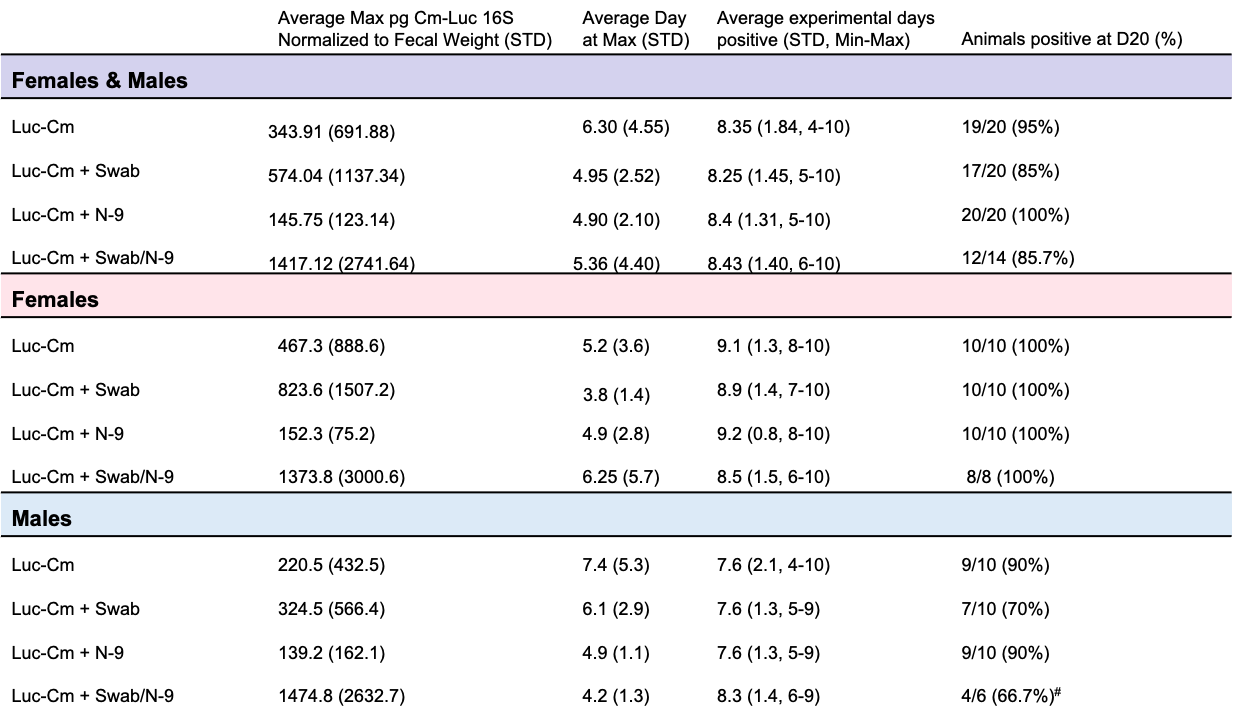

**Supplementary Table 1. Time course of Luc-Cm fecal shedding.** Feces from BALB/c mice (n=20/group, male and female) were collected on days 0-20 after anorectal infection with Luc-Cm. Feces were process to extract DNA and RealTime-qPCR was used to determine the pg of Luc-Cm 16S normalized to fecal weight using the formula (C × VT)/(VU × M). The average max pg Luc-Cm 16S normalized to fecal weight, average day at max pg Luc-Cm 16S normalized to fecal weight, average number of experiment days that were positive (> 1 pg of Luc-Cm 16S), and animals positive at day 20 were described for all of the groups, with either the sexes combined or separated by sex. Statistical analysis was performed utilizing a non-parametric Kruskal-Wallis One-Way ANOVA except for animals positive at day 20, which was analyzed using a Fisher’s exact test. The different epithelial disruption groups were compared to the No Disruption group. * *p* < 0.05, ** *p* < 0.01, *** *p* < 0.001, **** *p* < 0.0001. # 1 male removed, did not pooping on the final day.

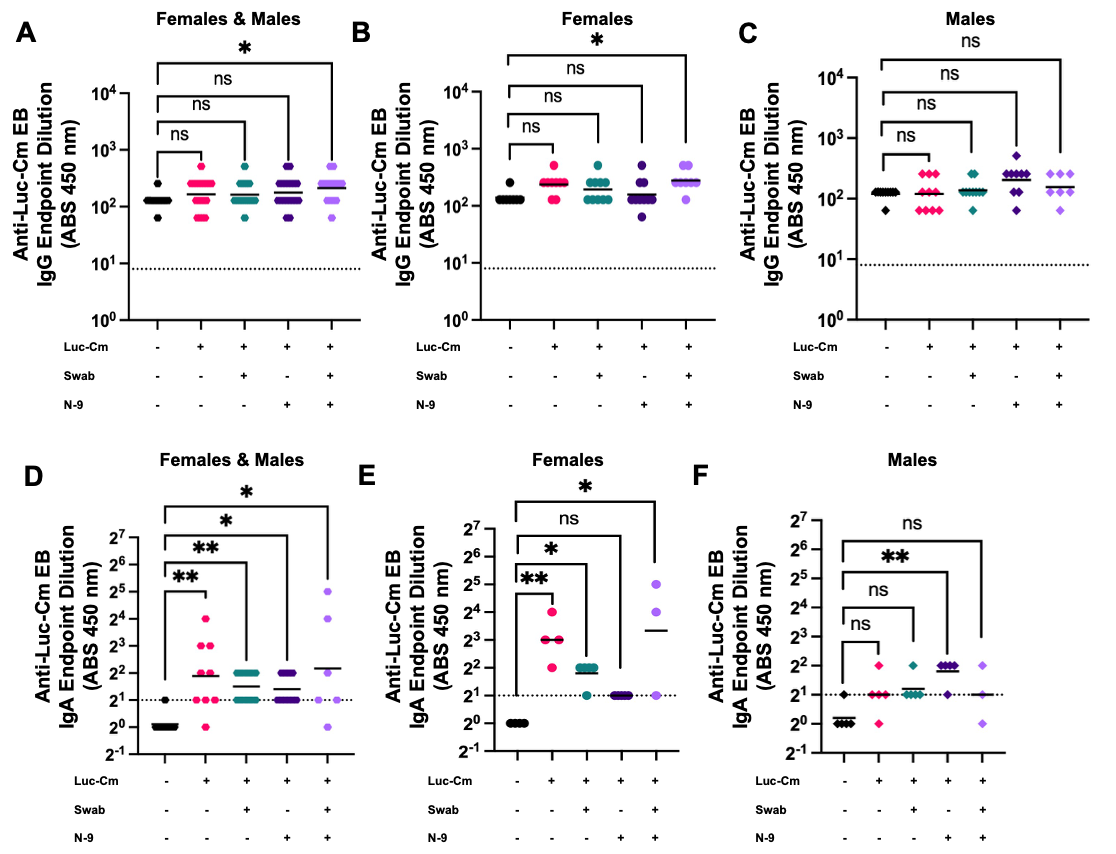

**Supplementary Figure 3. Antibody binding to Luc-Cm EBs post-infection.** On day 20 post-infection, antibody binding capacity to Luc-Cm EBs were determined using ELISA. The sera IgG bind capacity is shown as endpoint dilution titers for (**A**) females and males combined, (**B**) females, and (**C**) males. The sera IgA bind capacity is shown as endpoint dilution titers for (**D**) females and males combined, (**E**) females, and (**F**) males. Statistical analysis was performed utilizing a non-parametric Kruskal-Wallis One-Way ANOVA. ns = non-significant, * *p* < 0.05, ** *p* < 0.01, *** *p* < 0.001, **** *p* < 0.0001.

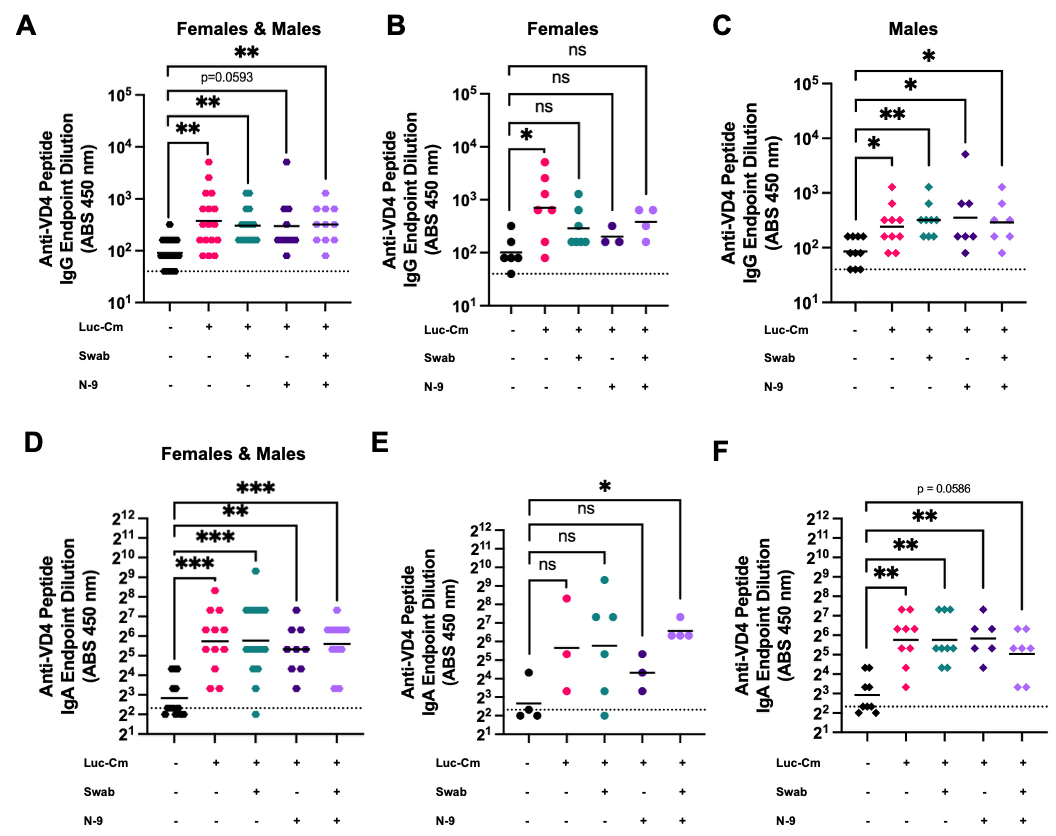

**Supplementary Figure 4: Antibody binding to VD4 peptide post-infection.** On day 20 post-infection, antibody binding capacity to Cm VD4 peptide were determined using ELISA. The sera IgG bind capacity was shown as endpoint dilution titers for (**A**) females and males combined, (**B**) females, and (**C**) males. The sera IgA bind capacity was shown as endpoint dilution titers for (**D**) females and males combined, (**E**) females, and (**F**) males. Statistical analysis was performed utilizing a non-parametric Kruskal-Wallis One-Way ANOVA. ns = non-significant, * *p* < 0.05, ** *p* < 0.01, *** *p* < 0.001, **** *p* < 0.0001.

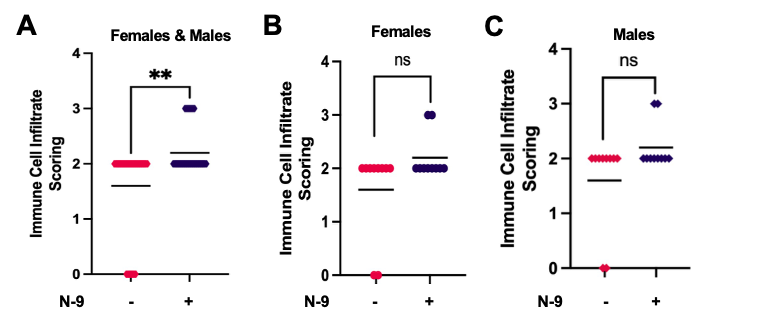

**Supplementary Figure 5:** BALB/c mice (n=20) mock vaccinations with PBS 3-times, 3-weeks apart. On day 68, the mice received either no disruption (n=20) or chemical disruption (n=20) treatment in the anorectal tract. The next day fecal material was collected, the mice were euthanized, and the distal colon was collected for histology. The severity and extent of inflammatory cell infiltration of the distal colon that were stained with H&E was determine using blinded scoring by a pathologist for either the sexes combined (**A**) or separated in females (**B**) and males (**C**). Statistical analysis was performed utilizing a non-parametric Mann-Whitney t-test. ns = non-significant, * *p* < 0.05, ** *p* < 0.01, *** *p* < 0.001, **** *p* < 0.0001.
